# The Integrative Conjugative Element (ICE) of *Mycoplasma agalactiae*: key elements involved in horizontal dissemination and influence of co-resident ICEs

**DOI:** 10.1101/308072

**Authors:** Eric Baranowski, Emilie Dordet-Frisoni, Eveline Sagné, Marie-Claude Hygonenq, Gabriela Pretre, Stéphane Claverol, Laura Fernandez, Laurent Xavier Nouvel, Christine Citti

**Affiliations:** IHAP, Université de Toulouse, INRA, ENVT, Toulouse, France; Pôle Protéomique, Centre de Génomique Fonctionnelle, Université de Bordeaux, Bordeaux, France

## Abstract

The discovery of integrative conjugative elements (ICEs) in wall-less mycoplasmas and the demonstration of their role in massive gene flows within and across species has shed new light on the evolution of these minimal bacteria. Of these, ICEA of the ruminant pathogen *Mycoplasma agalactiae* represents a prototype and belongs to a new clade of the Mutator-like superfamily that has no preferential insertion site and often occurs as multiple chromosomal copies. Here, functional genomics and mating experiments were combined to address ICEA functions and define the minimal ICEA chassis conferring conjugative properties to *M. agalactiae*. Data further indicated a complex interaction among co-resident ICEAs, since the minimal ICEA structure was influenced by the occurrence of additional ICEA copies that can *trans*-complement conjugative-deficient ICEAs. However, this cooperative behavior was limited to the CDS14 surface lipoprotein, which is constitutively expressed by co-resident ICEAs, and did not extend to other ICEA proteins including the *cis*-acting DDE recombinase and components of the mating channel whose expression was only sporadically detected. Remarkably, conjugative-deficient mutants containing a single ICEA copy knocked-out in *cdsl4* can be complemented by neighboring cells expressing CDS14. This result, together with the conservation of CDS14 functions in closely related species, may suggest a way for mycoplasma ICEs to extend their interaction outside of their chromosomal environment. Overall, this study provides a first model of conjugative transfer in mycoplasmas and offers valuable insights towards the understanding of horizontal gene transfer in this highly adaptive and diverse group of minimal bacteria.

**IMPRTANCE:** Integrative conjugative elements (ICEs) are self-transmissible mobile genetic elements that are key mediators of horizontal gene flow in bacteria. Recently, a new category of ICEs has been identified that confer conjugative properties to mycoplasmas, a highly adaptive and diverse group of wall-less bacteria with reduced genomes. Unlike classical ICEs, these mobile elements have no preferential insertion specificity and multiple mycoplasma ICE copies can be found randomly integrated into the host chromosome. Here, the prototype ICE of *Mycoplasma agalactiae* was used to define the minimal conjugative machinery and propose the first model of ICE transfer in mycoplasmas. This model unveils the complex interactions taking place among co-resident ICEs and suggests a way for these elements to extend their influence outside of their chromosomal environment. These data pave the way for future studies aiming at deciphering chromosomal transfer, an unconventional mechanism of DNA swapping that has been recently associated with mycoplasma ICEs.

## INTRODUCTION

Integrative conjugative elements (ICEs) are self-transmissible mobile genetic elements that are key mediators of horizontal gene flow in bacteria (1). These self-transmissible elements encode their excision, transfer by conjugation and integration into the genome of the recipient cell where they replicate as a part of the host chromosome. Recently, a new family of self-transmissible integrative elements has been identified in the genome of several mycoplasma species that confers conjugative properties to this important group of bacteria (2–8).

Mycoplasmas are well-known for having some of the smallest genomes thus far characterized in free-living organisms, with many species being successful human and animal pathogens (9, 10). Mycoplasmas belong to the class Mollicutes, a large group of atypical bacteria that have evolved from low GC, Gram-positive common ancestors (11). For decades, their evolution has been considered as marked by a degenerative process, with successive losses of genetic material resulting in current mycoplasmas having no cell wall and limited metabolic capacities (9). The recent discovery of massive horizontal gene transfer (HGT) in mycoplasmas has shed new light on the dynamics of their reduced genomes (12, 13). Evidence for HGT in these minimal bacteria came from the identification of putative ICEs in several species together with *in silico* data suggesting that mycoplasma species of distant phylogenetic groups have exchanged a significant amount of chromosomal DNA (14). Conjugative properties of mycoplasmas were further demonstrated using the ruminant pathogen *Mycoplasma agalactiae* as a model organism (7, 12). In this species, mating experiments and associated next generation sequencing established that mycoplasma ICEs (MICEs) are self-transmissible mobile elements conferring the recipient cells with the capacity to conjugate (Fig. 1). These uncovered at the same time an unconventional conjugative mechanism of chromosomal transfers (CTs), which involved large chromosomal regions and were independent of their genomic locations (12). While ICE self-dissemination was documented from ICE-positive to ICE-negative cells, CTs were observed in the opposite direction resulting in the incorporation of large genomic regions (Fig. 1). Remarkably, CTs can mobilize up to 10 % of the mycoplasma genome in a single conjugative event generating a complex progeny of chimeric genomes that may resemble conjugative distributive transfers in *Mycobacterium smegmatis* (15). While ICE and CTs appeared as two independent events, CTs rely on ICE factors most likely for providing the conjugative pore.

**FIG 1.**
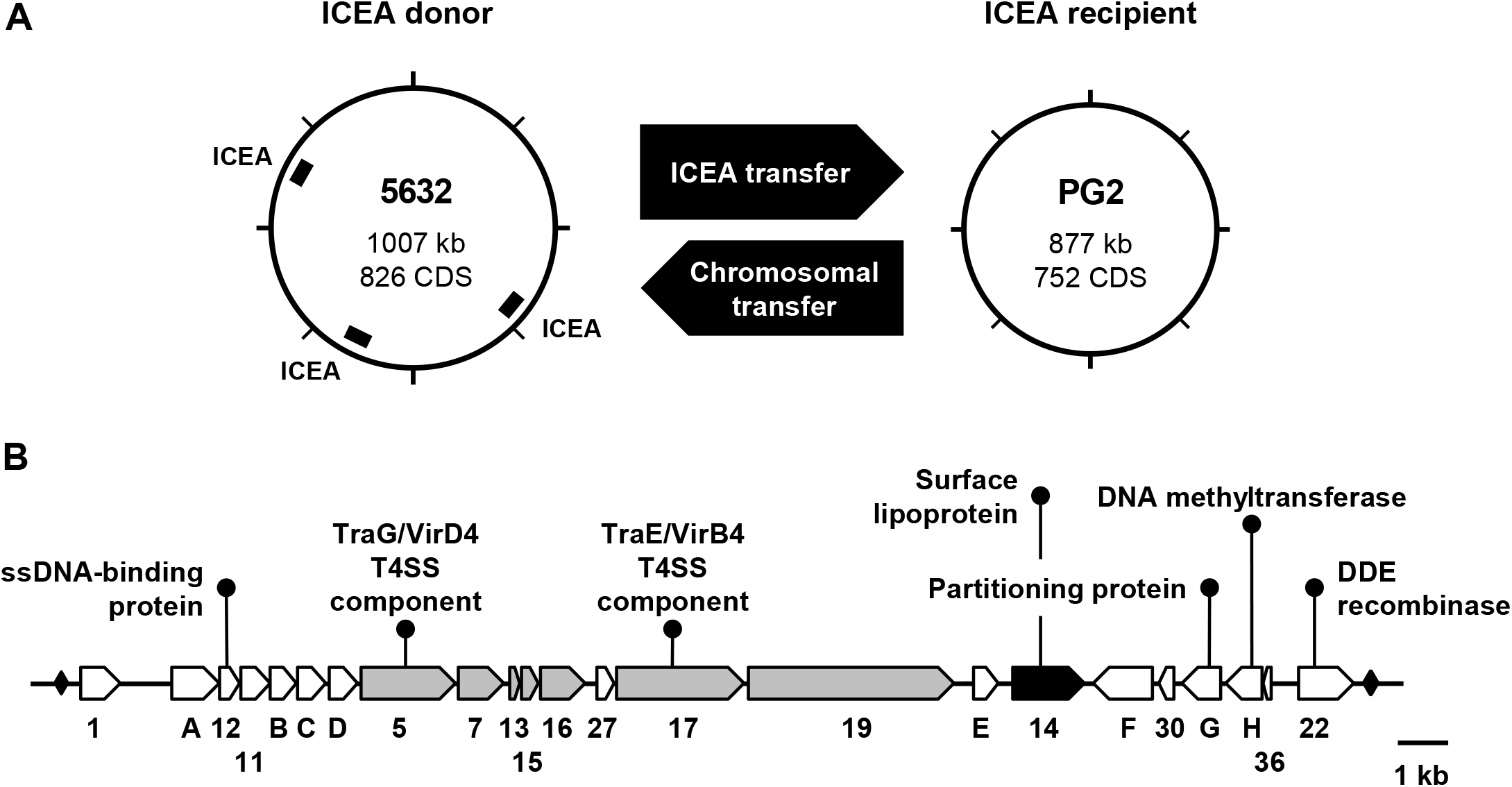
ICEA mediated horizontal gene transfers (HGT) in *M. agalactiae*. Schematic illustrating the two mechanisms of gene exchanges occurring upon mating experiments involving strain 5632 as ICE donor and strain PG2 as ICE recipient cells (7, 12) **(A)**. One of the three chromosomal ICEA copies of 5632 is transferred to PG2 and integrates randomly in the recipient genome (ICEA transfer). ICEA self-dissemination is associated with a second mechanism of gene exchange that occurs in the opposite direction from the recipient to the donor cells and involves large chromosomal DNA movements (Chromosomal transfer). ICEA transfer confers conjugative properties to the PG2 recipient cells (7). The 23 genes identified in ICEA are represented with their respective orientation and approximate nucleotide size **(B)**. The two inverted repeats (IR) flanking the ICEA are represented by black diamonds. The genes encoding predicted surface lipoproteins or proteins with putative transmembrane domains are in black and grey, respectively. Hypothetical functions were deduced from putative conserved domains found in several ICEA products (Table S1).

Of the MICEs so far described, the ICE of *M. agalactiae* (ICEA) has been most extensively studied (3, 16). ICEA and MICEs in general belong to a new family of self-transmissible integrative elements that rely on a DDE transposase of the prokaryotic Mutator-like family for their mobility (7, 17). Mainly associated with small and simple transposons such as insertion sequences, DDE transposases are also encoded by more complex mobile elements, such as streptococci Tn*GBS* conjugative transposons (17). Unlike Tn*GBS* that have a preferential insertion upstream of σA promoters, ICEA integration occurs randomly in the host chromosome generating a diverse population of ICEA-transconjugants (7, 17). This situation also contrasts with more conventional ICEs, which encode site-specific tyrosine recombinases (1). ICEA occurrence varies among *M. agalactiae* strains with strain 5632 containing three nearly identical ICEA copies and PG2 that contains no ICE, or a vestigial form (14, 16). Functional ICEAs are about 27 kb long and are composed of 23 genes (Fig. 1), most of which encode proteins of unknown function (Table S1) with no homolog outside of the Mollicutes (3). Among the few exceptions are CDS5 and CDS17, two proteins with similarity to conjugation-related TraG/VirD4 and TraE/VirB4, respectively (Table S1). Both proteins are energetic components of the type IV secretion systems, which are usually involved in DNA transport (18).

The establishment of laboratory conditions in which ICE transfer can be reproduced and analyzed in *M. agalactiae* (7), together with the development of specific genetic tools for the manipulation of this species (19), offer a unique opportunity to further investigate the detailed mechanisms underlying HGTs in mycoplasmas. In the present study, a transposon-based strategy was devised to knock-out individual ICEA genes and to decipher ICEA functions in *M. agalactiae*. Data showed that the minimal ICEA chassis required for conferring conjugative properties to *M. agalactiae* was influenced by the occurrence of additional ICEA copies that can *trans*-complement conjugative-deficient ICEAs. Complementation studies further unveil the complexity of this interplay that can even extend to neighboring cells and the key role played by the co-resident ICEA expression pattern. This study is a first step towards understanding HGT in mollicutes and provides a valuable experimental framework to decipher the mechanisms of DNA exchange in more complex bacteria when associated with this new category of mobile elements.

## RESULTS

### Conjugative properties of *Mycoplasma agalactiae* mutated ICEA

To elucidate the molecular mechanisms underlying ICE conjugative transfer in *M. agalactiae*, a library of 1,440 individual mutants was generated by random insertion of a mini-transposon (mTn) in the genome of strain 5632 that contains three nearly identical copies of a functional ICEA (Fig. 1). Mating experiments were conducted using pools of 96 individual 5632-mutants as donors and a pool of five PG2 recipient clones to avoid possible bias associated with a particular variant. Donors and recipients were chosen to carry compatible antibiotic markers (see Material and Methods) and resulting transconjugants were obtained with a frequency ranging from 2 × 10^−9^ to 8 × 10^−8^ transconjugants/total CFUs, as expected for 1:10 ratio (5632:PG2) that favors ICEA transfer from 5632 to PG2 (7). Double-resistant colonies were further subjected to detailed genetic analysis to (i) identify ICEA-positive PG2 versus 5632 having acquired PG2 genomic materials by CTs, and (ii) map the mTn position within ICEA-positive PG2 transconjugants. This strategy allowed the identification of 27 unique mutant ICEAs (Fig. 2A and Table S2). Remarkably, mTn insertions were found to cluster within a 6.4-kb ICEA region spanning *cdsE* to *cdsH* with the exception of three inserted in the noncoding regions *ncr1/A* and *ncr36/22* (Fig. 2A, mutants 3, 5 and 47), and one inserted in *cds11* (Fig. 2A, mutant 7).

**FIG 2.**
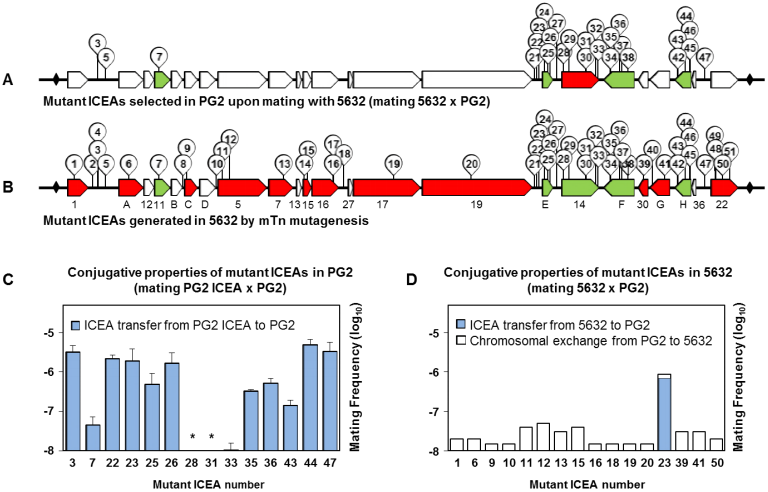
Functional analysis of mutant ICEAs in the 5632 and PG2 genetic backgrounds. Schematic illustrating the 51 mutant ICEAs generated by transposon mutagenesis in *M. agalactiae* strain 5632 **(B)** and the mutant ICEAs selected in PG2 upon mating with 5632 **(A)**. Individual mutant ICEAs are designated by their reference number together with the position of the mTn insertion (Table S2). The genes with no mTn insertion are indicated in white. ICEA genes found essential (red) or dispensable (green) are identified according to their genetic backgrounds that differ in their ICEA content (Fig. 1). Conjugative properties of selected mutant ICEAs in PG2 (**C**) and 5632 (**D**). Mating frequencies were calculated as the number of dual-resistant transconjugants per total CFUs (mating frequencies per single-resistant CFUs are provided in Table S3). Donor cells were mated with a pool of 5 ICEA-negative PG2 clones encoding resistance to puromycin (mating PG2 ICEA × PG2) or tetracycline (mating 5632 × PG2). Dual-resistant colonies were selected by using a combination of gentamicin and puromycin (mating PG2 ICEA × PG2), or gentamicin and tetracycline (mating 5632 × PG2). For matings PG2 ICEA × PG2 (**C**), the data represent means of at least three independent assays with the exception of mutant ICEA number 23 (9 independent assays). Since mutant ICEAs can be found integrated at different genomic positions, two PG2 ICEA transconjugants were used for mutant ICEA number 7 (ICEA at genomic position 395291 and 433901). Standard deviations are indicated by error bars. The asterisk indicates a mating frequency below the detection limit (1 × 10^−10^ transconjugants per total CFUs). For matings 5632 × PG2 (**D**), the data represent the average of two independent assays. The genetic profile of the transconjugants was determined using 10 to 166 dual-resistant colonies per mating, which for lower mating frequencies represent nearly all the progeny.

Since PG2 transconjugants contain no co-resident ICEA copies (Fig. S1), mating experiments were performed to evaluate the conjugative properties of selected mutant ICEAs (Fig. 2A). Individual PG2 transconjugants (further designated as PG2 ICEA) were mated with a pool of five ICEA-negative PG2 clones as recipient cells (Fig. 2C). The PG2 ICEA cells carrying a mTn inserted in *ncr1/A, ncr19/E* and *ncr36/22* (Fig. 2A and 2C, mutants 3, 22, 23 and 47) displayed comparable mating frequencies (1.9 to 3.5 × 10^−6^ transconjugants/total CFUs) suggesting that mTn insertions in these regions had no or minimal effect on conjugation. Conversely, mating experiments involving *cds14* knock-out ICEAs (Fig. 2A and 2C, mutants 28, 31, and 33) as in PG2 ICEA donor cells confirmed the essential role previously recognized for this gene (7), and further indicated that *cds14* can be complemented *in trans* by coresident ICEA copies, such as in 5632. The insertion of a mTn in *cds11, cdsE, cdsF* and *cdsH* (Fig. 2A and 2C, mutants 7, 25-26, 35-36, and 43-44) did not abrogate ICEA transfer, but several mutant ICEAs displayed a reduced capacity to self-disseminate. Whether the conjugative properties of these PG2 ICEA cells may be influenced by the chromosomal position of the integrated ICEA is unknown. However, similar mating frequencies (4 to 5 × 10^−8^ transconjugants/total CFUs) were observed for two PG2 transconjugants sharing the same mutant ICEA (Fig. 2C, mutant 7) integrated at different chromosomal sites (genomic positions 395291 and 433901). These results identified *cdsE, cdsF, cdsH*, and to a lesser extend *cds11*, as dispensable for ICEA self-dissemination.

### The minimal ICE chassis that confers conjugative properties to *Mycoplasma agalactiae*

As shown above, mutant ICEs recovered in PG2 transconjugants displayed a biased distribution of their mTn insertions (Fig. 2A). This raised the question of the representativity of the 5632 library and thus a PCR-based screening strategy for the direct identification of mutant ICEAs in 5632 was developed: mTn insertions across the entire ICEA were searched by a series of PCR assays using one primer matching each end of the mTn and one specific-ICEA primer selected from a set of oligonucleotides spanning the whole ICEA region. Amplifications were performed using pools of 96 individual mutants until one mTn insertion event per gene was detected, and positive pools were further characterized down to the single-mutant level. For each mutant, the mTn insertion was mapped by genomic DNA sequencing that also confirmed the presence of a single mTn per chromosome. Finally, the distribution of mTn insertions among the three ICEA copies of 5632 was determined by long-range PCR amplifications using mTn specific primers and a panel of oligonucleotides that are complementary to genomic DNA regions surrounding each ICEA copy. This strategy led us to identify 35 unique mutants (Table S2) among the three ICEA copies of 5632 (ICEA-I 29%; ICEA-II 37%; ICEA-III 34%). This time, mTn insertions were found broadly distributed throughout the entire ICEA locus with the exception of several genes all characterized by a small size ranging from 0.20 to 0.65 kb (*cds12, cdsB, cdsD, cds13, cds27* and *cds36*). These findings indicate that the particular set of mutant ICEAs selected above in PG2 cannot be simply explained by a poor representativity of the 5632 mutant library.

The 51 mutant ICEAs identified in 5632, either by PCR screening or by mating experiments, are illustrated in Figure 2B. Out of 35 mutant ICEAs identified by PCR, 24 did not correspond to detectable PG2 transconjugants previously obtained (compare Fig. 2A and 2B) suggesting that these mutant ICEAs have lost their capacity to disseminate from 5632 to PG2. This was confirmed by mating using individually each of these 5632 mutants as ICEA donor (Fig. 2D) and by further analyses of their progeny. Results showed that when transconjugants were obtained all displayed the 5632 genomic backbone of the mutant and correspond to 5632 having acquired the second PG2 antibiotic marker upon CTs. This was true for all but for 5632 mutant 23 (mTn inserted in *ncr19/E* with no influence on conjugation) that was used as a positive control for ICEA transfer and generated up to 97% of PG2 transconjugants (Fig. 2D, mutant 23).

Overall, these results indicate that ICEA transfer can be abrogated or strongly affected by disrupting the genes encoding CDSs 1, A, C, 5, 7, 15, 16, 17, 19, 30, G or 22 in 5632 (Fig. 2B). Unlike *cds14* knock-out ICEAs, the conjugative properties of these mutant ICEAs cannot be restored by co-resident ICEA copies. Finally, ICEA transfer was also abrogated by mTn insertion in *ncrD/5* and *ncr16/27* (Fig. 2B, mutants 10 and 18) raising questions about the presence of regulatory and/or *cis*-acting elements (*e.g., oriT*) in these regions. Whether short genes (< 0.65 kb) with no mTn insertion may encode essential functions remains to be further investigated.

### ICEA transfer in *Mycoplasma agalactiae* requires the CDS14 surface lipoprotein

The CDS14 lipoprotein is essential for mycoplasma conjugation and contains a 27 aa signal sequence (Fig. S2) that is characteristic of surface exposed lipoproteins in *M. agalactiae*. CDS14 surface location was confirmed in ICEA-positive cells by colony blotting assays using a specific antiserum (Fig. 3A) and is in agreement with proteomic data showing an association of CDS14 with the Triton X-114 hydrophobic fraction obtained after mycoplasma partitioning (16). Western blot analyses of 5632[ICEA cds14::mTn]^G^28 (mutant 28 in Fig. 2B and Table S2) having one of the three ICEA copies with a knock-out *cds14* further demonstrated that this lipoprotein can be expressed by co-resident ICEAs (Fig. 3B). This explains the capacity of *cdsl4* knock-out ICEAs to be horizontally transferred from 5632 cells observed above.

**FIG 3.**
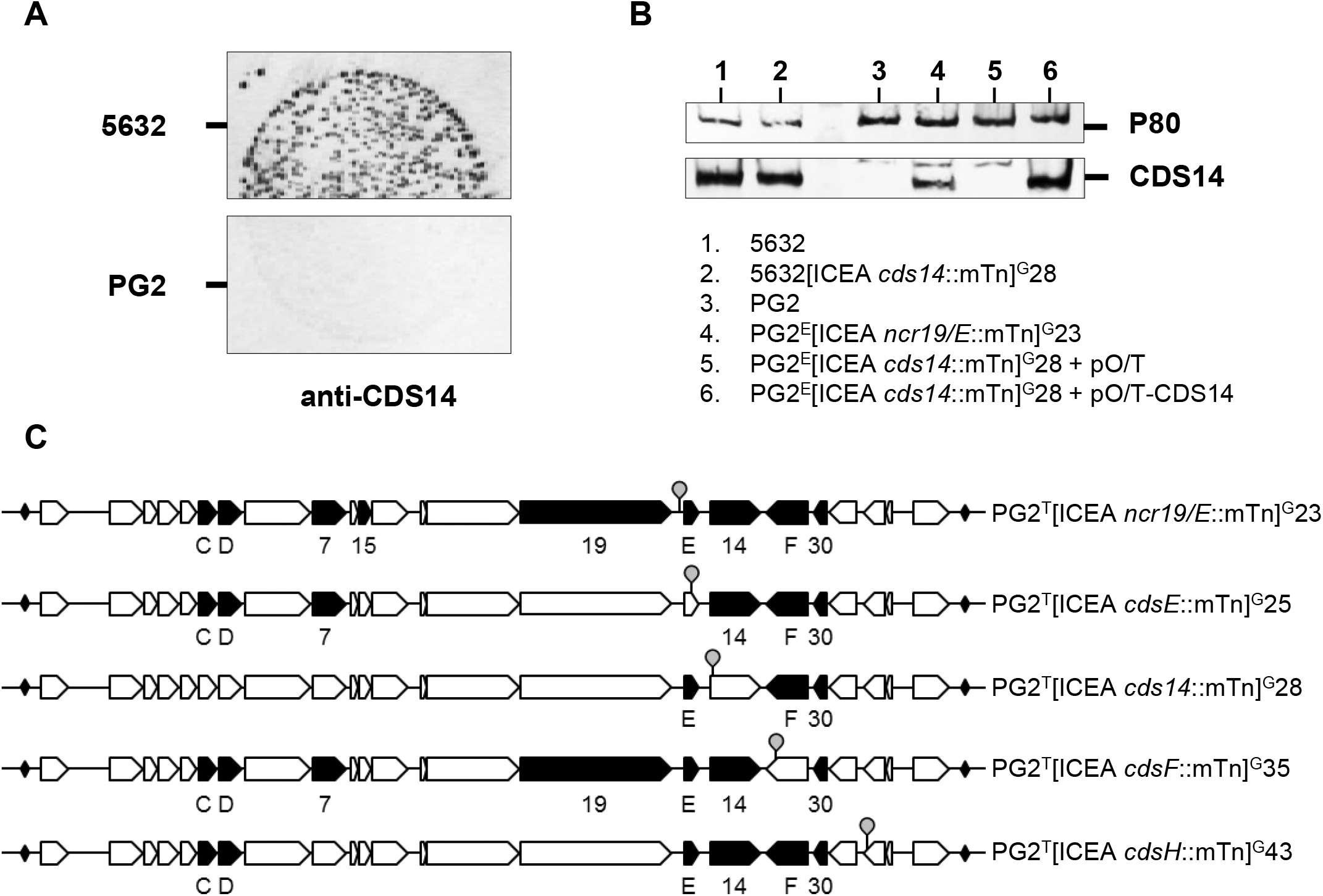
Protein expression of PG2-ICEA mutants. Immunostaining of *M. agalactiae* colonies showing CDS14 lipoprotein expression at the surface of 5632 cells (**A**). Colony blots were carried out by using a specific serum (anti-CDS14) and ICEA negative PG2 cells (PG2) were used as a negative control. Western blot analysis of CDS14 lipoprotein expression in 5632 and PG2 ICEA cells (**B**). CDS14 lipoprotein expression in 5632 that contains three chromosomal ICEA copies (1) was not abrogated in a 5632 mutant harboring a *cds14* knockout ICEA copy (2). CDS14 lipoprotein expression can be detected in PG2 transconjugants having acquired a mutant ICEA harboring a mTn inserted in *ncr19/E* (4), but not in PG2 (3) or PG2 transconjugants harboring a *cds14* knock-out ICEA (5). Transformation of PG2 transconjugants harboring a *cds14* knock-out ICEA with a plasmid expressing CDS14 restored the expression of the lipoprotein (6). A specific serum raised against lipoprotein P80 was used as control (P80). Schematic illustrating the protein expression profile of selected mutant ICEAs in PG2 cells (**C**). Mutant ICEAs are identified by their reference number (Table S2) and ICEA products detected by proteomics (Table S4) are indicated (closed arrows).

The role of the CDS14 lipoprotein was further investigated by using the conjugative-deficient PG2^e^[ICEA *cds14*::mTn]^G^28 having a only one ICEA copy with a mTn inserted in *cds14*. Complementation studies confirmed that the conjugative properties of this mutant can be restored upon transformation with plasmid pO/T-CDS14 that expresses the wild-type CDS14, but not with the empty vector (Table 1, matings A and B). Remarkably, transformation of ICE-negative recipient cells with pO/T-CDS14 also restored the conjugative properties of PG2^e^[ICEA *cds14*::mTn]^G^28 (Table 1, mating C) with only a reduction in mating frequency (ca. 20-fold). This result provides the first evidence of ICE complementation by neighboring cells and suggests that CDS14 lipoprotein may initiate ICEA transfer in *M. agalactiae* by promoting a contact between the donor and the recipient cells.

**Table 1.**
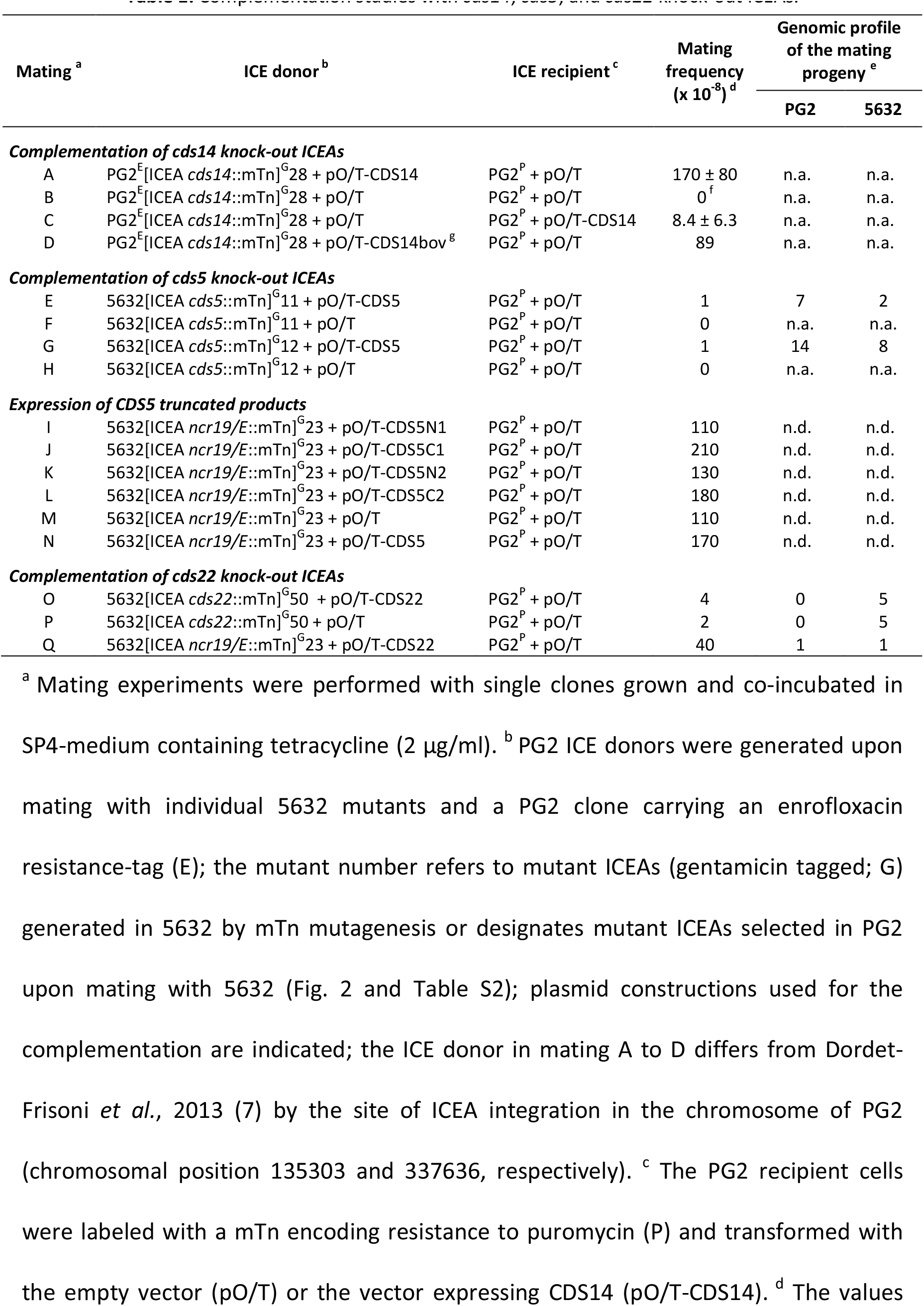
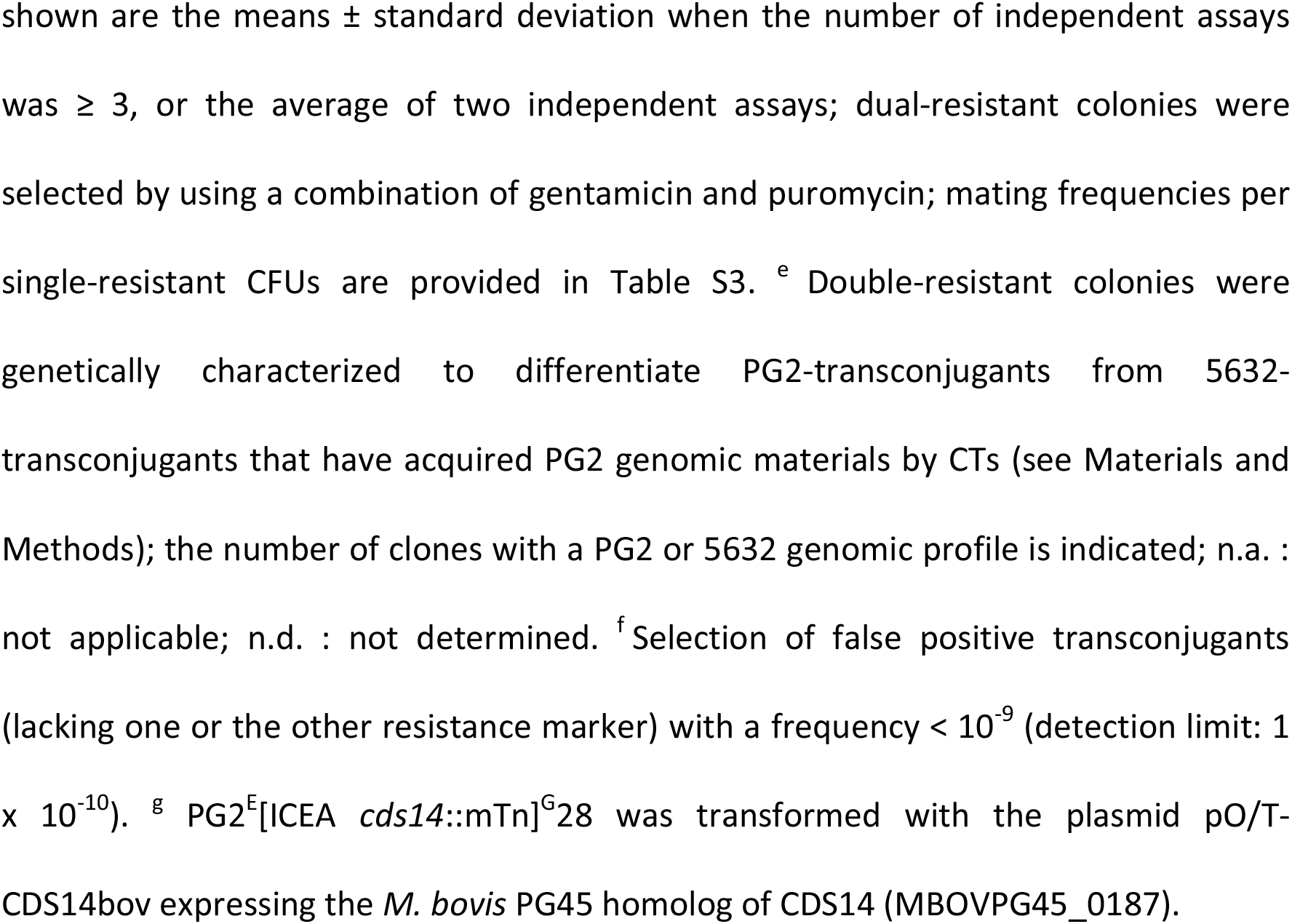
Complementation studies with *cds14, cds5*, and *cds22* knock-out ICEAs.

| Mating <sup>a</sup> | ICE donor <sup>b</sup> | ICE recipient <sup>c</sup> | Mating frequency (x 10 <sup>-8</sup>) <sup>d</sup> | Genomic profile of the mating progeny <sup>e</sup> |  |
| --- | --- | --- | --- | --- | --- |
|  |  |  |  | PG2 | 5632 |
| **Complementation of *cds14* knock-out ICEAs** |  |  |  |  |  |
| A | PG2<sup>E</sup>[ICEA *cds14*::mTn]<sup>G</sup>28 + pO/T-CDS14 | PG2<sup>P</sup> + pO/T | 170 ± 80 | n.a. | n.a. |
| B | PG2<sup>E</sup>[ICEA *cds14*::mTn]<sup>G</sup>28 + pO/T | PG2<sup>P</sup> + pO/T | 0<sup>f</sup> | n.a. | n.a. |
| C | PG2<sup>E</sup>[ICEA *cds14*::mTn]<sup>G</sup>28 + pO/T | PG2<sup>P</sup> + pO/T-CDS14 | 8.4 ± 6.3 | n.a. | n.a. |
| D | PG2<sup>E</sup>[ICEA *cds14*::mTn]<sup>G</sup>28 + pO/T-CDS14bov<sup>g</sup> | PG2<sup>P</sup> + pO/T | 89 | n.a. | n.a. |
| **Complementation of *cds5* knock-out ICEAs** |  |  |  |  |  |
| E | 5632[ICEA *cds5*::mTn]<sup>G</sup>11 + pO/T-CDS5 | PG2<sup>P</sup> + pO/T | 1 | 7 | 2 |
| F | 5632[ICEA *cds5*::mTn]<sup>G</sup>11 + pO/T | PG2<sup>P</sup> + pO/T | 0 | n.a. | n.a. |
| G | 5632[ICEA *cds5*::mTn]<sup>G</sup>12 + pO/T-CDS5 | PG2<sup>P</sup> + pO/T | 1 | 14 | 8 |
| H | 5632[ICEA *cds5*::mTn]<sup>G</sup>12 + pO/T | PG2<sup>P</sup> + pO/T | 0 | n.a. | n.a. |
| **Expression of *CDS5* truncated products** |  |  |  |  |  |
| I | 5632[ICEA *ncr19/E*::mTn]<sup>G</sup>23 + pO/T-CDS5N1 | PG2<sup>P</sup> + pO/T | 110 | n.d. | n.d. |
| J | 5632[ICEA *ncr19/E*::mTn]<sup>G</sup>23 + pO/T-CDS5C1 | PG2<sup>P</sup> + pO/T | 210 | n.d. | n.d. |
| K | 5632[ICEA *ncr19/E*::mTn]<sup>G</sup>23 + pO/T-CDS5N2 | PG2<sup>P</sup> + pO/T | 130 | n.d. | n.d. |
| L | 5632[ICEA *ncr19/E*::mTn]<sup>G</sup>23 + pO/T-CDS5C2 | PG2<sup>P</sup> + pO/T | 180 | n.d. | n.d. |
| M | 5632[ICEA *ncr19/E*::mTn]<sup>G</sup>23 + pO/T | PG2<sup>P</sup> + pO/T | 110 | n.d. | n.d. |
| N | 5632[ICEA *ncr19/E*::mTn]<sup>G</sup>23 + pO/T-CDS5 | PG2<sup>P</sup> + pO/T | 170 | n.d. | n.d. |
| **Complementation of *cds22* knock-out ICEAs** |  |  |  |  |  |
| O | 5632[ICEA *cds22*::mTn]<sup>G</sup>50 + pO/T-CDS22 | PG2<sup>P</sup> + pO/T | 4 | 0 | 5 |
| P | 5632[ICEA *cds22*::mTn]<sup>G</sup>50 + pO/T | PG2<sup>P</sup> + pO/T | 2 | 0 | 5 |
| Q | 5632[ICEA *ncr19/E*::mTn]<sup>G</sup>23 + pO/T-CDS22 | PG2<sup>P</sup> + pO/T | 40 | 1 | 1 |

<sup>a</sup> Mating experiments were performed with single clones grown and co-incubated in
SP4-medium containing tetracycline (2 µg/ml). <sup>b</sup> PG2 ICE donors were generated upon
mating with individual 5632 mutants and a PG2 clone carrying an enrofloxacin
resistance-tag (E); the mutant number refers to mutant ICEAs (gentamicin tagged; G)
generated in 5632 by mTn mutagenesis or designates mutant ICEAs selected in PG2
upon mating with 5632 (Fig. 2 and Table S2); plasmid constructions used for the
complementation are indicated; the ICE donor in mating A to D differs from Dordet-
Frisoni *et al.*, 2013 (7) by the site of ICEA integration in the chromosome of PG2
(chromosomal position 135303 and 337636, respectively). <sup>c</sup> The PG2 recipient cells
were labeled with a mTn encoding resistance to puromycin (P) and transformed with

Interestingly, global alignment of CDS14 lipoprotein with its homologs found in ICEB, the conjugative element occurring in the closely related *M. bovis* species, revealed 86.9% of sequence similarity (Fig. S2). To test whether these differences may influence the conjugative transfer of ICEA in *M. agalactiae*, PG2^E^[ICEA *cds14*::mTn]^G^28 was transformed with plasmid pO/T-CDS14bov expressing the ICEB CDS14 lipoprotein. This plasmid was able to restore the conjugative properties of *cds14* knock-out ICEA with a 2-fold reduction in mating frequency (Table 1, mating D). This suggests that one of the CDS14 functions is conserved between ICEA and ICEB.

Altogether these results unveiled the critical role played by ICE encoded surface lipoproteins in the exchange of genetic information within mycoplasma species and most likely across species.

### CDS5 expression from co-resident ICEAs is a key factor for *cds5* knock-out ICEA complementation

In contrast to *cds14* mutants that can be complemented by co-resident ICEAs, or even by neighboring cells expressing CDS14, a large number of mutant ICEAs were unable to disseminate from 5632 to PG2 (Fig. 2 and Table S2). Several of these mutant ICEAs were knocked-out in genes whose products were not detected by proteomics (see below) raising the question of whether the level of gene expression in co-resident ICEAs can influence their cooperative behavior. To address this issue, complementation studies were performed using plasmid DNA constructs expressing either CDS5 or CDS22, two proteins suspected to be involved in very distinct steps of ICE transfer.

While *cds5* knock-out ICEAs have lost their capacity to disseminate from 5632 to PG2 (Fig. 2D, mutants 11 and 12), mating of 5632 complemented with *cds5* (plasmid pO/T-CDS5) with ICEA-negative PG2 recipient cells resulted in a ca. 10-fold increase in mating frequency (Table 1, matings E and G). Analysis of the mating progeny revealed that 64 to 78% of these transconjugants displayed a PG2 genomic profile, indicating the conjugative transfer of the *cds5* knock-out ICEA from 5632 to PG2. Transformation with the empty vector (plasmid pO/T) as negative control resulted in no detectable event of transfer (Table 1, matings F and H). These data showed that *cds5* knock-out ICEAs can be complemented *in trans*, at least partially, when *cds5* is expressed from the expression vector, while paradoxically co-resident ICEAs were unable to restore the conjugative properties of *cds5* knock-out ICEAs. Finally, these complementation studies allowed us to rule out any lethal effect resulting from *cds5* knock-out ICEAs integration in the PG2 chromosome.

The CDS5 is a putative membrane bound hexamer with an ATPase activity displaying some similarity with the TraG/VirD4 conjugative channel component found in more classical bacteria (3). The formation of a hexameric structure by *cds5* products remains to be confirmed, but this multimeric organization may provide an alternative scenario for the inactive *cds5* knock-out ICEAs in 5632. Indeed, mTn insertion in *cds5* could lead to the expression of truncated products interfering with the hexamer complex formation, and thus inducing a negative dominant effect on co-resident ICEAs. To address this issue, truncated versions of *cds5* were cloned into the pO/T expression vector leading to plasmids pO/T-CDS5 N1, N2, C1, and C2, expressing respectively CDS5 N- and C-terminal regions resulting from mTn insertion in *cds5* mutants 11 and 12 (Fig. S3). Transformation of 5632[ICEA *ncr19/E*::mTn]^G^23 (mTn inserted in *ncr19/E* with no influence on conjugation) with constructions carrying truncated forms of *cds5*, the full-length *cds5* or the empty vector had no influence on mating efficacy (Table 1, matings I to N), indicating that the expression of CDS5 truncated products did not inhibit ICEA transfer, and that the conjugative-deficient phenotype of *cds5* knock-out mutants is not the result of a negative dominant effect.

### CDS22 expression is unable to *trans-complement cds22* knock-out ICEAs

A second series of complementation studies were performed with the *cds22* knock-out ICEA mutant 5632[ICEA *cds22*::mTn]^G^50. This gene encodes a DDE recombinase that was previously shown to mediate ICEA excision and circularization (7). Transformation with pO/T-CDS22 did not increase mating frequency when compared to the empty vector, and no PG2 transconjugants were identified upon analysis of the mating progeny (Table 1, matings O and P). Transformation of 5632[ICEA *ncr19/E*::mTn]^G^23 (mTn inserted in *ncr19/E* with no influence on conjugation) with pO/T-CDS22 or the empty vector had no or minimal influence on the mating frequencies (Table 1, matings Q and M). These results suggest that the *cds22* knockout ICEA cannot be complemented *in trans*, neither by co-resident ICEAs nor by a CDS22 expressing plasmid. This result is consistent with the longstanding observation that DDE transposases show a *cis*-preference for their activities (20–22).

Altogether these results illustrate the complex interactions taking place among co-resident ICEAs in 5632, and elucidated some of the mechanisms underlying their non-cooperative behavior.

### Protein expression profiles of PG2-ICEA mutants

A previous study has shown that in 5632 three ICEA products are detectable by proteomic analysis under laboratory conditions: CDS14, and to a lower extent CDS17 and CDS30 (16). To further characterize ICEA expression in different genomic contexts, a proteomic analysis was conducted using a set of PG2 transconjugants having acquired a mutated ICEA copy from 5632 (Fig. 3C and Table S4). Data revealed that up to 9 ICEA products, namely CDSs C, D, 7, 15, 19, E, 14, F, and 30, were detected in PG2^T^[ICEA *ncr19/E*::mTn]^G^23 (mTn inserted in *ncr19/E* with no influence on conjugation). CDS17 was also detected in PG2^T^[ICEA *ncr19/E*::mTn]^G^23 but below cut-off values. CDSE, CDS14 and CDSF were not detected in PG2^T^[ICEA *cdsE*::mTn]^G^25, PG2^T^[ICEA *cds14*::mTn]^G^28 or PG2^T^[ICEA *cdsF*::mTn]^G^35, in which the corresponding genes are disrupted. Besides confirming the disruption of these genes, these data also indicate that mTn insertion in *cdsE* has no polar effect on the expression of the downstream *cds14*. Interestingly, the data suggested that some ICEA loci might be downregulated to undetectable levels in the conjugative-deficient PG2^T^[ICEA *cds14*::mTn]^G^28. These corresponded to CDSC, CDSD, CDS7 and CDS19 detected in other mutants, whose genes are located upstream of *cds14*. Overall, PG2 ICEA mutants that were tested here and disrupted in identified coding genes, versus *ncr19/E*, had a simplified ICEA protein expression profile.

## DISCUSSION

Since their discovery in mycoplasma species of the Hominis phylogenetic group, MICEs have been found broadly distributed across Mollicutes (2–6, 8) and their pivotal role in HGTs is emerging (7, 12, 13). Taking advantage of the *M. agalactiae* ICE prototype, this study provides the first functional analysis of MICE factors involved in conjugative transfer. Because MICEs, such as ICEA, are often found in multiple copies, this study points toward their complex interplay in the mycoplasma host-environment.

### The functional ICEA backbone

The minimal ICEA chassis conferring conjugative properties to *M. agalactiae* was identified by random transposon mutagenesis. Of the 23 genes reported in ICEA, 17 were found disrupted by the insertion of a mTn and 13 were found essential for self-dissemination, since a single mTn insertion in any of these regions abrogated the conjugative properties of *M. agalactiae*. These data point towards the minimal ICEA machinery being composed of (i) a cluster of 7 proteins with predicted transmembrane domains that most likely represents a module associated with the conjugative channel (CDS5 to CDS19), (ii) a surface exposed lipoprotein (CDS14), (iii) a putative partitioning protein (CDSG), (iv) a DDE transposase (CDS22), and (v) several other proteins with no predicted function (CDS1, CDSA, CDSC, and CDS30). The conjugative-deficient phenotype of the ICEA mutants is unlikely to be result of a polar effect since (i) mTn insertions were identified at close proximity to essential ICEA regions with no influence on conjugation *(cds1, cds14, cds30*, and to a lesser extent *cdsA* and *cdsG)*, (ii) *cds14* knock-out ICEAs can be complemented *in trans*, (iii) ICEA-mutants having the putative channel module disrupted by a mTn inserted in *cds5* can be plasmid-complemented, and (iv) mTn insertions in the *cdsE-cdsH* region have no influence on protein expression from surrounding genes. Whether additional essential ICEA functions may be encoded by several of the 6 short genes (0.20 to 0.65 kb) with no mTn insertion remains to be further investigated.

The minimal ICEA chassis was consistent with the conservation of *cds5, cds17, cds19* and *cds22* across documented ICEs of ruminant mycoplasma species (8), and the occurrence of *cds1, cds14* and *cds16* at very similar location in a majority of MICEs (2, 4–6, 23). Interestingly, the conjugative properties of *M. agalactiae* were also abrogated by mTn insertion in NCRs raising questions regarding the presence of regulatory elements and/or key motifs, such as an *oriT*, in these regions. The occurrence of such sequences in the NCR1/A (1238 nucleotides) is supported by the identification of a hairpin motif (TGGCTCAT-N_5_-ATGAGCCA) at positions 2046 to 2066 (S. Torres-Puig, personal communication). Whether mTn insertion in NCRs may influence the expression surrounding ICEA regions is unknown.

Accessory ICEA functions were only found associated with 4 genes, namely *cds11, cdsE, cdsF* and *cdsH*. These accessory functions are all encoded within a 6.4-kb ICEA region spanning *cdsE* to *cdsH*, with the exception of *cds11* that belongs to a cluster of 6 genes *(cdsA* to *cdsD)* located upstream the putative channel module (Fig. 1). Although dispensable, an important reduction of the mating frequency was observed for several mutants having a mTn inserted in these ICEA regions (Fig. 2C). For PG2^T^[ICEA *cds11*::mTn]^G^7 (Fig. 2A, mutant 7), this reduction was not influenced by the position of the mutant ICEA in the PG2 chromosome. Finally, Blastp analyses with CDSE revealed a significant similarity (≥ 90%) to a putative prophage gene product found in the chromosome of PG2 (MAG6440) and 5632 (MAGa7400). The presence of this chromosomal *cdsE* homolog is puzzling and its role in ICEA transfer remains to be confirmed.

### A minimal genome but coping with multiple ICEA copies

Unlike classical ICEs, ICEA has no preferential insertion specificity and multiple copies can be found at different loci of the host chromosome. This raised questions regarding their maintenance in the small mycoplasma genomes and the deleterious effect that can be associated with their random insertion, in particular because they were found within coding sequences (7, 16). Whether ICEA may confer any advantage *in vivo* is unknown, but PG2 ICEA transconjugants displayed a reduced fitness in laboratory conditions (unpublished data). Many bacterial ICEs and some prokaryotic transposable elements carry cargo genes implicated in accessory functions, such as antibiotic resistance, which confer a selective advantage to their host (1). Such cargo genes have never been reported in MICEs, but ICE mediated CTs are likely contributing to the acquisition of new phenotypic traits upon chromosomal exchanges.

### Backup functions associated with co-resident ICEAs

Our data suggest that co-resident ICEAs are able to cooperate by complementing essential functions in mutant ICEAs. This was shown by using *cds14* knock-out ICEAs, which can self-disseminate when occurring in the context of 5632, but not in PG2 that contains no additional ICEA copy. The complementation of *cds14* by co-resident ICEAs was further confirmed by the constitutive expression of the CDS14 lipoprotein in 5632 mutants having one of the 3 ICEA copy knocked-out in *cds14*, but not in PG2 cells having acquired a *cds14* knock-out ICEA. Remarkably, this cooperative behavior was found to extend to neighboring cells, since transformants of ICEA-negative cells containing a plasmid vector expressing CDS14 were able to complement *cds14* knock-out ICEAs in neighboring cells. Besides providing the first example of ICE complementation by neighboring cells, this result has deep implications for the dissemination of MICEs within and across mycoplasma species. This was further supported by complementation studies showing that the CDS14 lipoprotein in *M. agalactiae* can be substituted by its homolog in *M. bovis* ICEB, and our previous study showing ICEA mediated CTs between *M. agalactiae* and *M. bovis* (12).

Interestingly, the cooperative behavior documented with *cds14* knock-out ICEAs did not extend to other critical ICEA regions. Complementation studies with *cds22* knock-out ICEAs confirmed that several critical ICEA functions can be associated with cis-acting elements that cannot be complemented by co-resident ICEAs. However, studies with *cds5* mutant ICEAs suggested that interactions among co-resident ICEAs can be more complex. Indeed, *cds5* knock-out ICEAs can be *trans*-complemented upon transformation with a CDS5 expressing plasmid but not by co-resident ICEAs. Since ICEA transfer from 5632 to PG2 occurs only at low frequency, ICEA activation is expected to be a rare event. It is thus reasonable to speculate that only one of the three chromosomal ICEA copies in 5632 can be stochastically activated. This hypothesis provides a simple scenario to understand the lack of complementation of *cds5* mutants by co-resident ICEAs, since this component of the mating channel is expected to be only expressed upon ICEA activation. It is further supported by proteomic analysis showing a simplified ICEA protein expression profile that contrasted with the constitutive expression of the CDS14 surface lipoprotein.

### Conclusions

The results generated in the present study were combined with current knowledge to propose the first working model of horizontal ICE dissemination in mycoplasmas, including cooperation among co-resident ICEs (Fig. 4). These data, together with the large collection of ICEA-mutants generated in this study, pave the way for future studies aiming at deciphering ICE-mediated CTs within and among mycoplasma species. These simple organisms also provide a valuable experimental frame to decipher the mechanisms of DNA exchange in more complex bacteria when associated with this new category of mobile elements.

**FIG 4.**
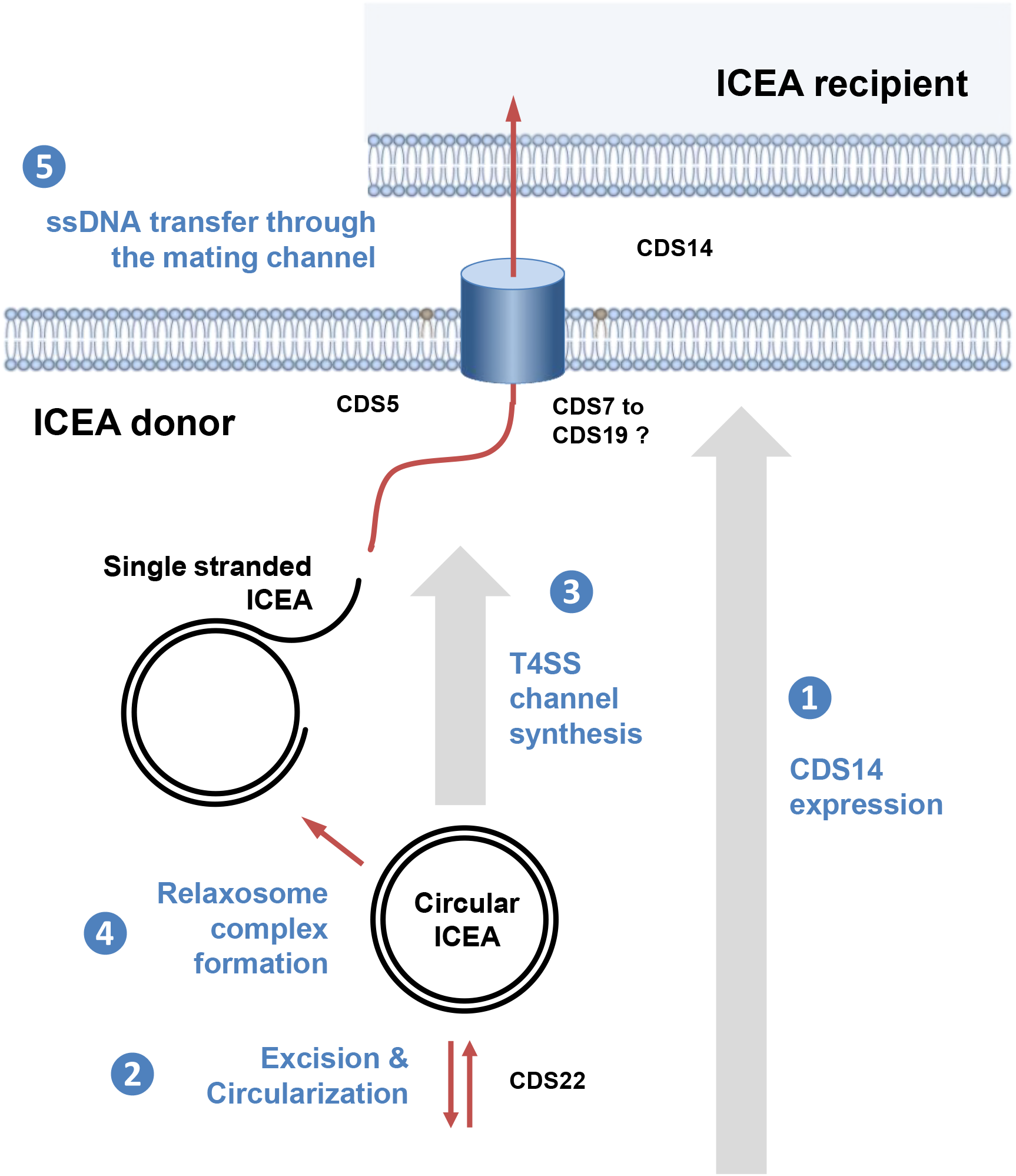
Overview of conjugative ICE transfer in *M. agalactiae*. This schematic was adapted from Alvarez-Martinez and Christie (18), and illustrates the 5 key steps in ICEA transfer. Upon normal conditions, ICEA copies are found integrated into the host chromosome and most ICEA genes are not expressed. Among the few proteins expressed by chromosomal ICEAs is the CDS14 lipoprotein that is surface exposed and plays a critical role in initiating the conjugative process (1). When ICEA gene expression is induced, by specific cellular conditions or stochastically, the *cis*-acting DDE transposase is produced and one of the three ICEA copies excises from the chromosome and forms a circular dsDNA molecule (2). ICEA circularization induces the expression of the conjugative module, whose products assemble into the mating pore, a simplified form of T4SS found in more complex bacteria (3). A protein complex, known as relaxosome, recognizes the origin of transfer *(oriT)* on the circular ICEA and a relaxase generates a linear ssDNA by nicking the ICEA DNA (4). Finally, the relaxosome complex interacts with the TraG-like (VirD4 homologue) energetic component found at the inner side of the membrane that facilitates the transfer of the ssDNA bound to the relaxase through the mating channel (5). Once in the recipient strain, the ICEA re-circularizes, becomes double stranded and integrates randomly into the host chromosome. The minimal functional ICEA encompasses 80% of the coding sequence and includes a gene cluster *(cds5* to *cds19)* encoding proteins with transmembrane domains that most likely represents a module associated with the conjugative channel. Additional essential ICEA determinants included the CDS14 surface lipoprotein, the CDSG putative partitioning protein and the DDE transposase (CDS22), together with several proteins of unknown functions (CDSs 1, A, C, and 30).

## MATERIALS AND METHODS

### Mycoplasmas and culture conditions

*M. agalactiae* strains PG2 and 5632 have been previously described (14, 16), and the sequence of each genome is available in databases (GenBank reference sequences CU179680.1 and FP671138.1, respectively). These two strains differs in their ICE content with strain 5632 having three almost identical chromosomal copies of ICEA (ICEA-I, -II, and -III), while strain PG2 contains only a severely degenerated, vestigial ICE (14, 16). *M. agalactiae* was grown at 37 °C in SP4 medium supplemented with 500 μg/ml cephalexin (Virbac). When needed, gentamicin (50 μg/ml), tetracycline (2 μg/ml) or puromycin (10 μg/ml), was added to the medium, alone or in combination. Due to their small cell- and colony-size, mycoplasma growth cannot be monitored by optical density. Mycoplasma titers were thus determined based on colonies counts on solid media after 4 to 7 days of incubation at 37° C, using a binocular stereoscopic microscope (19).

### Transposon mutagenesis and genetic tagging of mycoplasmas with antibiotic markers

A similar approach was used for transposon mutagenesis and genetic tagging of *M. agalactiae*. Selective antibiotic markers were introduced randomly in the mycoplasma genome as previously described by transforming mycoplasma cultures with the plasmid pMT85 or its derivatives (7, 19, 24). The pMT85 carries a mini-transposon (mTn) derived from the gentamicin resistance *Tn4001*. The transposase gene *(tnpA)* is located outside of the mTn to prevent re-excision events once it is inserted in the host chromosome (19). Two derivatives, pMT85-Tet and pMT85-Pur, were constructed as previously described by replacing the gentamicin resistance gene with a tetracycline or a puromycin resistance marker, respectively (7).

### PCR-based screening of mycoplasma mutant library

A set of 19 oligonucleotides spanning the whole ICEA region (Table S5) was used to develop a PCR-based screening of the mutant library and identify 5632 mutants having a mTn inserted within ICEA regions. Each ICEA specific primer was used in combination with the transposon-specific oligonucleotide SG5 priming at both inverted repeats (IRs) that define the extremities of the integrated transposon (Table S5). PCR amplifications were performed according to the recommendations of the Taq DNA polymerase supplier (New England Biolabs). For each mutant, the position of the mTn insertion in the *M. agalactiae* chromosome was determined by sequencing the junction between *M. agalactiae* genomic DNA and the 5’- or 3’-end of the transposon using oligonucleotides SG6_3pMT85E (specific to the 5’-end of the Gm-tagged version of the mTn), SG9pMM21-7mod (specific to the 5’-end of the Tet-tagged version of the mTn) or EB8 (specific to the 3’-end of all mTn constructions) as primers (Table S5). Genomic DNA sequencing was performed at the Genomic platform GeT-Purpan (Toulouse, France). The distribution of mTn insertions among the three ICEA copies was determined by long-range PCR amplifications (Expand Long Template PCR System; Roche Life Science) using mTn specific primers and a panel of oligonucleotides corresponding to genomic DNA regions surrounding each ICEA locus (Table S5).

### Mycoplasma mating experiments and genetic characterization of transconjugant progenies

Mating experiments were conducted as described previously by co-incubation of ICE-positive and ICE-negative cells (7). Mycoplasma growth may considerably vary from batch to batch when using the rich SP4 that contains serum and yeast extract. To reduce potential bias in mating frequencies observed in between experiments, a single batch of medium was used in this study. Cultures of donor and recipient mycoplasmas (10^9^ CFUs) were mixed in a 1:1 ratio (matings PG2 ICEA × PG2) or a 1:10 ratio (matings 5632 × PG2) to increase the chances of recovering PG2 ICEA transconjugants. The mating frequency was calculated by dividing the number of double-resistant colonies obtained on selective solid media by the number of mycoplasma colonies obtained on non-selective media. *M. agalactiae* transconjugants were characterized by PCR amplification using genomic DNA prepared from individual colonies (7). Presence of antibiotic resistance genes and ICEA in transconjugants was confirmed by using specific oligonucleotides (Table S5). The nature of the genetic backbone was addressed by using a set of primers pairs that covers the *M. agalactiae* genome and produces PCR fragments specific to 5632 or PG2 (Table S5), as previously described (7, 12).

### DNA constructs for protein expression in mycoplasmas

Protein expression in *M. agalactiae* was performed as previously described by using the plasmid p20-1miniO/T (designated in the present study as pO/T) (19, 25). Briefly, mycoplasma coding sequences were cloned downstream of the lipoprotein P40 gene (MAG2410) promoter region. These two regions were assembled by PCR amplification using overlapping primers (Table S5). The resulting PCR product was cloned into pGEM-T Easy (Promega) before subcloning at the *NotI* site of the pO/T. PCRs were performed using the Phusion high-fidelity DNA polymerase (New England Biolabs). DNA constructions were verified by DNA sequencing and introduced in *M. agalactiae* by transformation, as previously described (19).

### Proteomic analyses and immunodetection of ICEA products

*M. agalactiae* grown under normal and mating growth conditions were subjected to proteomic analyses. Cells were collected by centrifugation of mycoplasma cultures (8,000 × *g*), washed and resuspended in Dulbecco’s phosphate-buffered saline (DPBS). Proteins were separated by 1D SDS-PAGE and gel sections were subjected to trypsin digestion. Peptides were further analyzed by nano liquid chromatography coupled to a nanospray Q-Exactive hybrid quadruplole-Orbitrap mass spectrometer (Thermo Scientific). Peptides were identified as previously described by using a database consisting of *M. agalactiae* strain 5632 entries (26). ICEA products were detected by specific anti-sera on Western and colony blots (25, 27). Triton-X114 soluble proteins were extracted from *M. agalactiae* as previously described (28). The anti-CDS14 lipoprotein rabbit serum was produced by animal immunization with a recombinant CDS14 protein (pMAL™ Protein Fusion and Purification System; New England Biolabs). A sheep serum raised against the *M. agalactiae* surface antigen P80 was used as a control (25). Western and colony blots were developed by using swine anti-rabbit or rabbit anti-sheep immunoglobulin G conjugated to horseradish peroxidase (DAKO) and the 4-chloro-naphthol substrate or the SuperSignal West Dura Extended Duration Substrate (Thermo Scientific).

## SUPPLEMENTAL MATERIAL

**Table S1**. Relevant features of ICEA products.

**Table S2**. Mutant ICEAs generated in *M. agalactiae* strain 5632.

**Table S3**. Mating frequencies per single-resistant CFUs.

**Table S4**. Proteomic analysis of PG2 ICEA mutants.

**Table S5**. Oligonucleotides used in the present study.

**Figure S1**. Mutant ICEAs selected in PG2 occur as a single ICEA copy randomly integrated in the host chromosome.

**Figure S2**. Global alignment of CDS14 lipoproteins found in ICEs of *M. agalactiae* strain 5632 and *M. bovis* strain PG45.

**Figure S3**. Plasmid constructions carrying *cds5* or truncated versions of *cds5*.

## ACKNOWLEDGMENTS

This work was supported by grant ANR09MIE016 (MycXgene) from the French national funding research agency (ANR) and financial supports from INRA and ENVT. We thank Richard Herrmann and Sebastien Guiral for providing pMT85 and pMT85-Tet plasmid constructions. We also thank Philippe Giammarinaro for his help in producing the anti-P80 sheep serum, as well as Emilie Houssin and Abdel Touré for excellent technical assistance. Finally, we also thank Oscar Q Pich and Sergi Torres-Puig for helpful discussions.

## SUPPLEMENTARY FIGURE LEGENDS

**FIG S1. Mutant ICEAs selected in PG2 occurs as a single ICEA copy randomly integrated in the host chromosome.** Southern blotting experiments (E. Dordet-Frisoni, M. S. Marenda, E. Sagné, L. X. Nouvel, R. Guérillot, P. Glaser, A. Blanchard A, F. Tardy, P. Sirand-Pugnet, E. Baranowski, and C. Citti, Mol Microbiol 89:1226-1239, 2013, doi:10.1111/mmi.12341) with PG2 ICEA transconjugants failed to reveal multiple ICEA integrations. Mycoplasma genomic DNAs were restricted with *EcoRV* and hybridized with mTn Gm-specific (A) or ICEA *cds22-* specific (B) probes. The identification of a single Gm-positive DNA fragment in PG2 ICEA transconjugants was in agreement with genomic DNA sequencing data indicating a single mutant-ICEA insertion in the PG2 chromosome. This result was further supported by DNA hybridization with the cds22-specific probe that also discarded any wild-type ICEA copy in PG2 ICEA transconjugants. Differences in size between *cds22-*positive DNA fragments are consistent with the random insertion of ICEA in the host chromosome. The digested ICEA circular form is indicated by an arrow. For each PG2 ICEA transconjugants, the size of Gm positive DNA fragments was in agreement with predicted values (C). The number in parenthesis indicates the size of the ICEA fragment with an inserted mTn. The *Gm*- and *cds22*-positive fragments are indicated by asterisks (*: *Gm*-specific probe; **: *cds22*-specific probe). Dashed lines indicate a fragment overlapping ICEA and genomic DNA.

**FIG S2. Global alignment of CDS14 lipoproteins found in ICEs of *M. agalactiae* strain 5632 (5632) and *M. bovis* strain PG45 (PG45).** The alignment of CDS14 lipoprotein sequences derived from 5632 (MAGa5010) and PG45 (MBOVPG45_0187) were performed by using Needleman-Wunsch global alignment. The 27 aa sequence characteristic of surface exposed lipoproteins is underlined.

**FIG S3. Plasmid constructions carrying *cds5* or truncated versions of *cds5***. Schematics illustrating plasmid constructions carrying the full-length *cds5* (pO/T-CDS5), or truncated versions of *cds5* (pO/T-CDS5 N1, C1, N2 and C2). Truncated sequences are CDS5 N- and C-terminal regions resulting from mTn insertion in 5632[ICEA cds5::mTn]^G^11 and 5632[ICEA cds5::mTn]^G^12 (Fig. 2B and Table S2). Coding sequences were cloned downstream of the *M. agalactiae* lipoprotein P40 gene (MAG2410) promoter region (arrow).

## REFERENCES

1. Johnson CM, Grossman AD. 2015. Integrative and Conjugative Elements (ICEs): what they do and how they work. Annu Rev Genet 49:577–601.

2. Calcutt MJ, Lewis MS, Wise KS. 2002. Molecular genetic analysis of ICEF, an integrative conjugal element that is present as a repetitive sequence in the chromosome of Mycoplasma fermentans PG18. J Bacteriol 184:6929–6941.

3. Marenda M, Barbe V, Gourgues G, Mangenot S, Sagne E, Citti C. 2006. A new integrative conjugative element occurs in Mycoplasma agalactiae as chromosomal and free circular forms. J Bacteriol 188:4137–4141.

4. Pinto PM, Carvalho, MO, Alves-Junior, L, Brocchi, M, Schrank, IS. 2007. Molecular analysis of an integrative conjugative element, ICEH, present in the chromosome of different strains of Mycoplasma hyopneumoniae. Genet Mol Biol 30:256–263.

5. Wise KS, Calcutt MJ, Foecking MF, Röske K, Madupu R, Methé BA. 2011. Complete genome sequence of Mycoplasma bovis type strain PG45 (ATCC 25523). Infect Immun 79:982–983.

6. Thiaucourt F, Manso-Silvan L, Salah W, Barbe V, Vacherie B, Jacob D, Breton M, Dupuy V, Lomenech AM, Blanchard A, Sirand-Pugnet P. 2011. Mycoplasma mycoides, from “mycoides Small Colony” to “capri”. A microevolutionary perspective. BMC Genomics 12:114.

7. Dordet-Frisoni E, Marenda MS, Sagné E, Nouvel LX, Guérillot R, Glaser P, Blanchard A, Tardy F, Sirand-Pugnet P, Baranowski E, Citti C. 2013. ICEA of Mycoplasma agalactiae: a new family of self-transmissible integrative elements that confers conjugative properties to the recipient strain. Mol Microbiol 89:1226–1239.

8. Tardy F, Mick V, Dordet-Frisoni E, Marenda M, Sirand-Pugnet P, Blanchard A, Citti C. 2014. Integrative conjugative elements (ICEs) are widespread in field isolates of Mycoplasma species pathogenic for ruminants. Appl Environ Microbiol 81:1634–1643.

9. Razin S, Yogev D, Naot Y. 1998. Molecular biology and pathogenicity of mycoplasmas. Microbiol Mol Biol Rev 62:1094–1156.

10. Citti C, Blanchard A. 2013. Mycoplasmas and their host: emerging and re-emerging minimal pathogens. Trends Microbiol 21:196–203.

11. Woese CR, Maniloff J, Zablen LB. 1980. Phylogenetic analysis of the mycoplasmas. Proc Natl Acad Sci USA 77:494–498.

12. Dordet-Frisoni E, Sagné E, Baranowski E, Breton M, Nouvel LX, Blanchard A, Marenda MS, Tardy F, Sirand-Pugnet P, Citti C. 2014. Chromosomal transfers in mycoplasmas: when minimal genomes go mobile. MBio 5:e01958.

13. Citti C, Dordet-Frisoni E, Nouvel LX, Kuo CH, Baranowski E. 2018. Horizontal gene transfers in mycoplasmas (Mollicutes). Curr Issues Mol Biol 29:3–22.

14. Sirand-Pugnet P, Lartigue C, Marenda M, Jacob D, Barré A, Barbe V, Schenowitz C, Mangenot S, Couloux A, Segurens B, de Daruvar A, Blanchard A, Citti C. 2007. Being pathogenic, plastic, and sexual while living with a nearly minimal bacterial genome. PLOS Genet 3:e75.

15. Gray TA, Krywy JA, Harold J, Palumbo MJ, Derbyshire KM. 2013. Distributive conjugal transfer in mycobacteria generates progeny with meiotic-like genome-wide mosaicism, allowing mapping of a mating identity locus. PLOS Biol 11:e1001602.

16. Nouvel LX, Sirand-Pugnet P, Marenda MS, Sagné E, Barbe V, Mangenot S, Schenowitz C, Jacob D, Barré A, Claverol S, Blanchard A, Citti C. 2010. Comparative genomic and proteomic analyses of two Mycoplasma agalactiae strains: clues to the macro- and micro-events that are shaping mycoplasma diversity. BMC Genomics 11:86.

17. Guérillot R, Siguier P, Gourbeyre E, Chandler M, Glaser P. 2014. The diversity of prokaryotic DDE transposases of the mutator superfamily, insertion specificity, and association with conjugation machineries. Genome Biol Evol 6:260–272.

18. Alvarez-Martinez CE, Christie PJ. 2009. Biological diversity of prokaryotic type IV secretion systems. Microbiol Mol Biol Rev 73:775–808.

19. Baranowski E, Guiral S, Sagné E, Skapski A, Citti C. 2010. Critical role of dispensable genes in Mycoplasma agalactiae interaction with mammalian cells. Infect Immun 78:1542–1551.

20. Nagy Z, Chandler M. 2004. Regulation of transposition in bacteria. Res Microbiol 155:387–398.

21. Chandler M, Fayet O, Rousseau P, Ton Hoang B, Duval-Valentin G. 2015. Copy-out-paste-in transposition of IS911: a major transposition pathway. Microbiol Spectr 3:MDNA3-0031-2014.

22. Duval-Valentin G, Chandler M. 2011. Cotranslational control of DNA transposition: a window of opportunity. Mol Cell 44:989–996.

23. Shu H-W, Liu T-T, Chan H-I, Liu Y-M, Wu K-M, Shu H-Y, Tsai S-F, Hsiao K-J, Hu WS, Ng WV. 2012. Complexity of the Mycoplasma fermentans M64 genome and metabolic essentiality and diversity among mycoplasmas. PLOS One 7:e32940.

24. Zimmerman C-U, Herrmann R. 2005. Synthesis of a small, cysteine-rich, 29 amino acids long peptide in Mycoplasma pneumon/ae. FEMS Microbiol Lett 253:315–321.

25. Skapski A, Hygonenq M-C, Sagné E, Guiral S, Citti C, Baranowski E. 2011. Genome-scale analysis of Mycoplasma agalactiae loci involved in interaction with host cells. PLOS One 6:e25291.

26. Crouzet M, Claverol S, Lomenech A-M, Sénéchal CL, Costaglioli P, Barthe C, Garbay B, Bonneu M, Vilain S. 2017. Pseudomonas aerug/nosa cells attached to a surface display a typical proteome early as 20 minutes of incubation. PLOS One 12:e0180341.

27. Nouvel L-X, Marenda M, Sirand-Pugnet P, Sagné E, Glew M, Mangenot S, Barbe V, Barré A, Claverol S, Citti C. 2009. Occurrence, plasticity, and evolution of the vpma gene family, a genetic system devoted to high-frequency surface variation in *Mycoplasma agalactiae*. J Bacteriol 191:4111–4121.

28. Baranowski E, Bergonier D, Sagné E, Hygonenq M-C, Ronsin P, Berthelot X, Citti C. 2014. Experimental infections with Mycoplasma agalactiae identify key factors involved in host-colonization. PLOS One 9:e93970.

